# Measles virus exits human airway epithelia via infectious center sloughing

**DOI:** 10.1101/2021.03.09.434554

**Authors:** Camilla E. Hippee, Brajesh K. Singh, Andrew L. Thurman, Ashley L. Cooney, Alejandro A. Pezzulo, Roberto Cattaneo, Patrick L. Sinn

## Abstract

Measles virus (MeV) is the most contagious human virus, but we do not fully understand why. Unlike most respiratory viruses, MeV does not infect the airway epithelium immediately. MeV traverses the epithelium within immune cells that carry it to lymphatic organs where amplification occurs. Infected immune cells then synchronously deliver large amounts of virus to the airways. However, our understanding of MeV replication in airway epithelia is limited. To model it, we use well-differentiated primary cultures of human airway epithelial cells (HAE) from lung donors. In HAE, MeV spreads directly cell-to-cell forming infectious centers that grow for ∼3-5 days, are stable for a few days, and then disappear. Transepithelial electrical resistance remains intact during the entire course of HAE infection, thus we hypothesized that MeV infectious centers may slough off while preserving epithelial function. After documenting by confocal microscopy that infectious centers progressively detach from HAE, we recovered apical washes and separated cell-associated from cell-free virus by centrifugation. Virus titers were about 10 times higher in the cell-associated fraction than in the supernatant. In sloughed infectious centers, ciliary beating persisted and apoptotic markers were not readily detected, suggesting that they retain functional metabolism. Cell-associated MeV infected primary human monocyte-derived macrophages, modeling the first stage of infection in a new host. Single-cell RNA sequencing identified wound healing, cell growth, and cell differentiation as biological processes relevant for infectious center sloughing. 5-ethynyl-2’-deoxyuridine (EdU) staining located proliferating cells underneath infectious centers. Thus, cells located below infectious centers divide and differentiate to repair the extruded infected epithelial patch. As an extension of these studies, we postulate that expulsion of infectious centers through coughing and sneezing could contribute to MeV’s strikingly high reproductive number by allowing the virus to survive longer in the environment and by delivering a high infectious dose to the next host.

**AUTHOR SUMMARY:** Measles virus (MeV) is a respiratory pathogen that infects millions worldwide each year. Although sometimes mischaracterized as an innocuous childhood disease, measles remains a leading cause of death for children under five. MeV is the most contagious human virus and requires vaccination rates above 90% to maintain herd immunity. Global decreases in vaccination rates over the past ten years contributed to recent, widespread MeV outbreaks. We uncover here a novel mechanism by which MeV exits the human airways that may explain why it is much more contagious than other viruses. We document that infected cells containing cell-associated virus slough *en masse* from the airway epithelial sheet. These expelled infectious centers are metabolically active and can transmit infection to primary human monocyte-derived macrophages more efficiently than cell-free virus particles. Thus, cell-associated MeV can transmit host-to-host, a new paradigm for efficient respiratory virus transmission.

## INTRODUCTION

Despite the development of an effective vaccine for measles virus (MeV), measles persists in populations that have limited access to healthcare and is reemerging in populations that refuse vaccinations. MeV outbreaks were extensive in 2019, with 1,282 confirmed cases in the United States and more than 500,000 confirmed cases worldwide [1]. MeV is of particular concern because of its high transmission potential, measured by the basic reproduction number (R_0_). MeV has an estimated R_0_ value between 12 and 18, which suggests vaccination rates should exceed 92% to protect a community via herd immunity [2-4]. Cases of MeV are projected to rise due to postponed measles vaccination campaigns as healthcare infrastructures focus on COVID-19 cases [5].

The MeV replication cycle is fundamentally different from that of other respiratory viruses [6-8]. MeV enters the body through the upper airways and infects alveolar macrophages and dendritic cells that express its primary receptor, the signaling lymphocytic activation molecule (SLAM) [9]. These cells ferry the infection through the epithelial barrier and spread it to the local lymph nodes [10, 11]. Amplification of MeV in immune tissues sets the stage for synchronous, massive invasion of tissues expressing the MeV epithelial receptor, nectin-4 [12, 13]. This two-phase process contributes to the extremely contagious nature of MeV [14-19].

However, knowledge of the respiratory phase of MeV infection is limited. To model it, we use well-differentiated primary cultures of human airway epithelial cells (HAE) that are maintained at an air-liquid interface. Contrary to initial assumptions, we demonstrated that MeV enters HAE from the basolateral side, delivered by infected immune cells [20, 21]. MeV infection of HAE is minimally cytopathic. Epithelial integrity, as monitored by transepithelial electrical resistance, remains intact for weeks after inoculation; in addition, infected cells retain their columnar structure and lateral cytoskeletal interactions without forming visible syncytia [22]. Using a recombinant MeV expressing green fluorescent protein (GFP), we observed that cytosolic GFP rapidly flows from infected into adjacent cells. These results suggest the formation of pores along the lateral membrane of columnar epithelial cells and provide a route for direct cell-to-cell spread [22]. Furthermore, using a MeV expressing GFP linked to a component of its ribonucleocapsids (RNP), we observed movement of RNPs along the circumapical F-actin rings of newly infected cells, a strikingly rapid mechanism of horizontal trafficking between epithelial cells [23].

In spite of efficient spread between respiratory epithelial cells, apical budding is inefficient: MeV titers in apical washes *in vitro* and in bronchial alveolar lavages of macaques *in vivo* are lower than those of other respiratory viruses [11, 21, 24, 25]. On the other hand, recent studies of MeV spread suggest that cell-associated virus may have a significant role in host-to-host transmission. Specifically, respiratory droplets with the highest viral titers were recovered during intervals when a patient was coughing most frequently [26]. An association between coughing and high viral titer secretions was also observed in experimentally infected macaques [25]. A closer look at these secretions revealed that MeV-infected cells containing cell-associated virus were expelled during coughing and the titers of cell-associated and cell-free virus within the secretions were similar. In this study, we investigated how infectious MeV is released from HAE. We present evidence suggesting that sloughing of metabolically active infectious centers contributes to MeV’s strikingly high reproductive number.

## RESULTS

### Infectious centers dislodge from HAE as units

Using a MeV that expresses green fluorescent protein (MeV-GFP), we infected HAE (MOI = 1) from the basolateral surface and live-imaged infectious centers at low power over a period of 3 weeks (**Fig 1A**). During the first ∼5 days of infection, MeV spreads to surrounding cells, causing the infectious centers to grow in size. Around 7-10 days post-infection, infectious centers often “disappear” from the epithelial sheet. To understand their fate, we performed confocal microscopy at an early time point, day 3 (**Fig 1B and Movie S1**), and a late time point, day 21 (**Fig 1C and Movie S2**). In contrast to day 3, at day 21 the infectious center was dislodging from the epithelial layer. The cells of the infectious center remained clustered while detaching from uninfected epithelia, causing the infectious center to shed as a unit, as shown in the 3D reconstruction models (**Figs 1D, 1E, and S1**).

**Fig 1.**
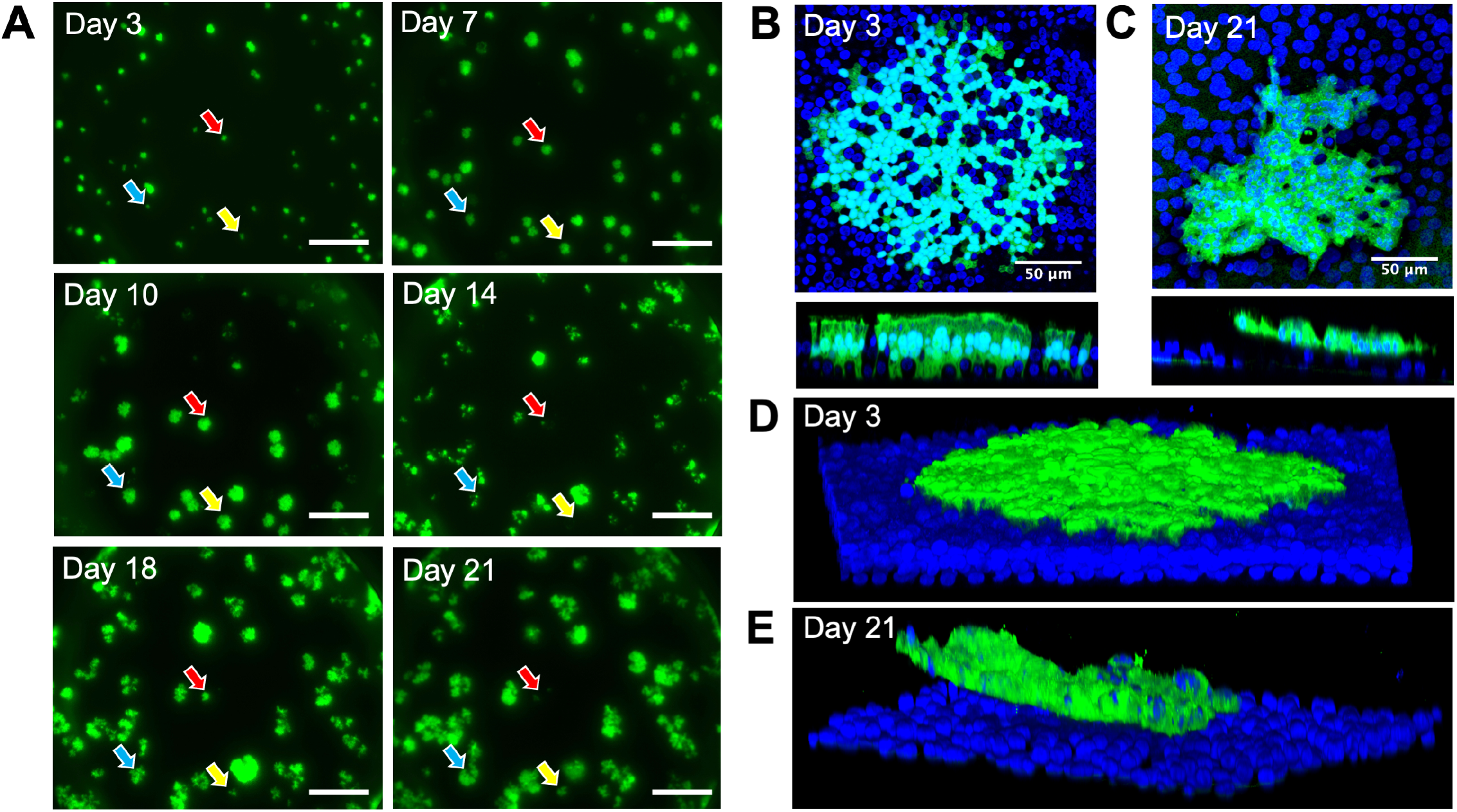
Infectious centers dislodge from HAE as units. (A) Live fluorescence microscopy of HAE infected with MeV-GFP (MOI = 1) over a time course of 21 days. All images are from the same field of view and are representative of 3 human donors. Colored arrows indicate examples of unique infectious centers that disappear during the time course. Scale bars = 500 μm. (B and C) *En face* and vertical confocal images of infectious centers at 3 days post-infection and 21 days post-infection, respectively. Z-stack images from B and C were used to create 3D models (D and E) respectively. Green, MeV-GFP; blue, DAPI.

### Sloughed infectious centers contain most released infectivity

To investigate the relevance of infectious center sloughing for virus transmission, we sought to quantify virus load in infected HAE cultures. We collected apical washes, cell lysates, and basolateral media from infected HAE every 3-4 days for 21 days post-infection. Apical washes were gently centrifuged in order to separate cell-free virus in the supernatant from cell-associated virus in the pellet (**Fig 2A**). We then measured virus titer in cell lysates, basolateral media, and cell-free and cell-associated virus from apical media (**Fig 2B**). High titers were observed in the cell lysates starting at 7 days post-inoculation, consistent with microscopy observations. In apical washes, virus titers were very low through day 10 post-inoculation. Starting from day 14, cell-associated virus titers were at least 10-fold higher than cell-free virus titers. These results indicate that most infectious MeV remains cell associated and exits the epithelial sheet via cell sloughing.

**Fig 2.**
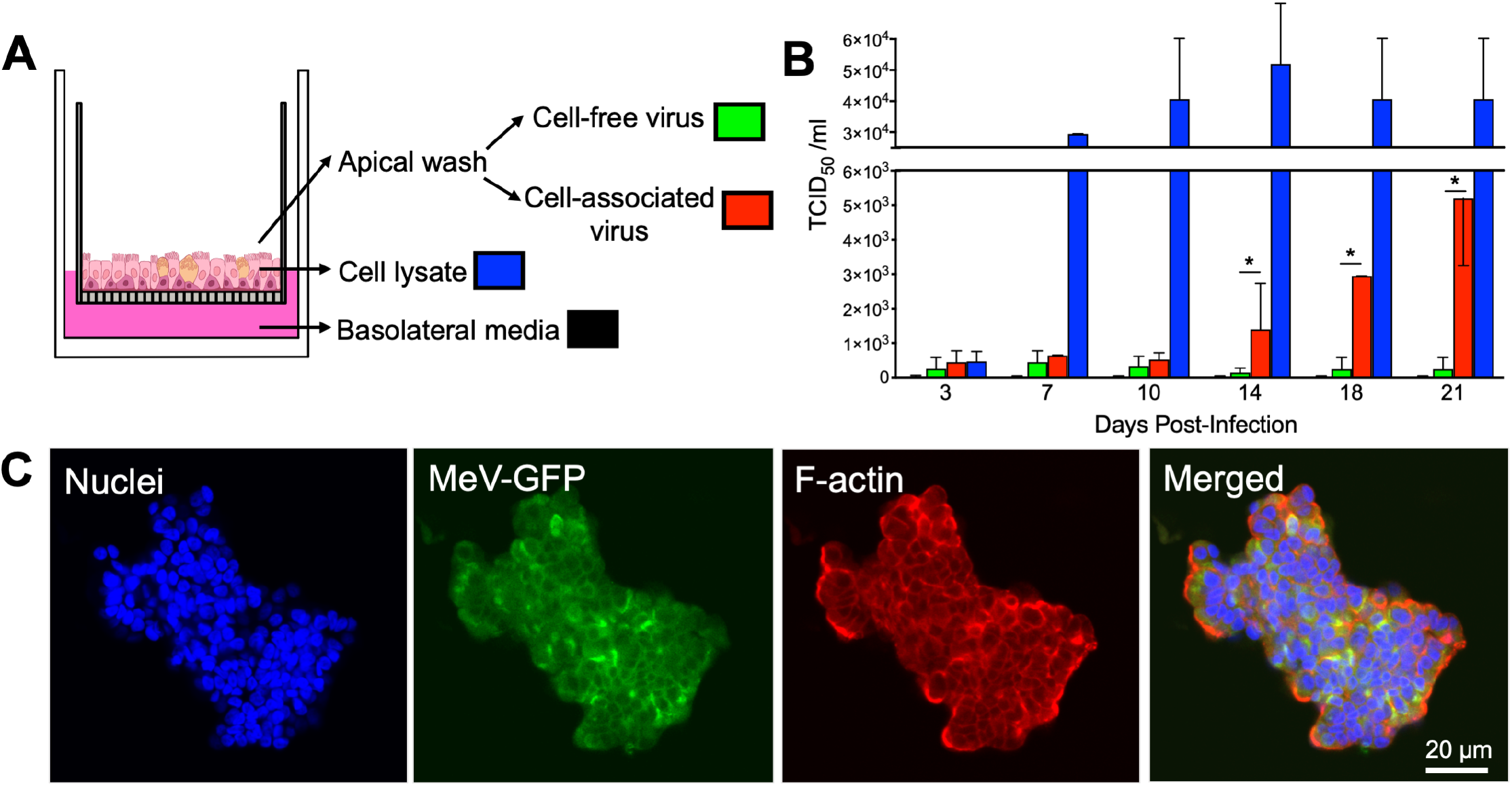
Sloughed infectious centers contain MeV. (A) Basolateral media, cell lysates, and apical washes were collected from HAE at 3, 7, 10, 14, 18, and 21 days post-infection (MOI = 1). Apical washes were gently centrifuged to separate cell-free virus from cell-associated virus. (B) TCID_50_ titers were performed on all four sample types at each timepoint (n = 3 human donors). Means ± standard deviation are shown. *p<0.05, cell-free vs. cell-associated. (C) Apical washes were mounted on coverslips and sloughed infectious centers were counterstained with DAPI (blue) and phalloidin (red). Images were collected with confocal microscopy and are representative of 3 human donors.

### Sloughed infectious centers remain viable

Infectious centers were collected in the apical washing to assess the viability after sloughing. Immunostaining and confocal microscopy imaging revealed intact nuclei and the F-actin cytoskeleton (**Fig 2C**). Strikingly, ciliary beating persisted in some sloughed infectious centers (**Movie S3**), which requires active metabolism [27].

To assess the extent to which viability is preserved in sloughed infectious centers, we used immunostaining to measure cleaved caspase-3, an apoptosis marker. Sloughed infectious centers were negative for caspase-3 staining **(Fig 3A)**; whereas, HAE treated with a positive control, protein kinase inhibitor staurosporine, were caspase-3 positive (**Fig 3B**). Western blotting confirmed that cleaved caspase-3 is not found in the lysates of mock or MeV-infected HAE over 14 days (**Fig 3C**). As an additional control, we used respiratory syncytial virus (RSV), another Paramyxovirus that induces apoptosis and apical cell sloughing in the bronchus of infants [28]. Caspase-3 and caspase-7 activity was significantly higher in RSV-infected HAE than in mock-infected HAE (**Fig 3D**), but these activities remained at background level in MeV-infected HAE. Altogether, these results indicate that the cells within MeV infectious centers remain viability after dislodging from the epithelial sheet.

**Fig 3.**
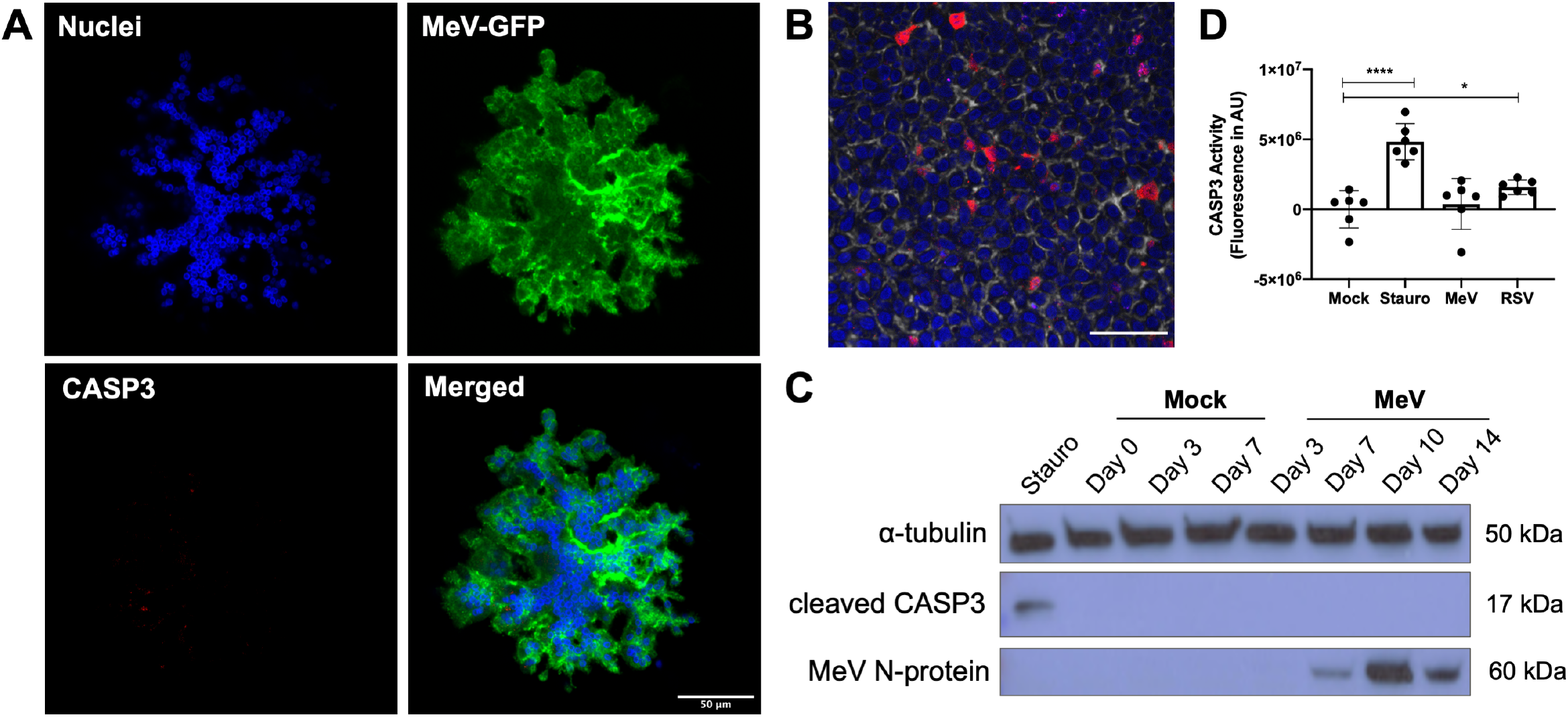
Cells of sloughed infectious centers are not apoptotic. (A) Apical washes were collected from MeV-infected HAE (14 days post-infection; MOI = 1), fixed, and immunostained for cleaved caspase-3 (CASP3). (B) HAE were treated with staurosporine (100 μM, 5 hrs) as a positive control to induce apoptosis, fixed, and immunostained for CASP3 (red), DAPI (blue), and phalloidin (gray). Scale bar = 50μm. (C) Western blot was performed on lysates from mock or MeV-infected HAE (n = 3; MOI = 1). Blots were probed for cleaved CASP3 and MeV N-protein. α-tubulin was used as a loading control protein. Staurosporine (stauro) treatment was used as a positive control. (D) Caspase-3 activity was assayed following mock, staurosporine (100 μM, 5 hrs), MeV (MOI = 1), or respiratory syncytial virus (RSV, MOI = 1) treatment of HAE (14 days post-infection; n = 3 human donors with 2 technical replicates). Fluorescence was measured in arbitrary units (AU) via plate reader. ****p < 0.0001; *p < 0.05.

### Sloughed infectious centers spread MeV infection to primary macrophages

We next asked if sloughed infectious centers infect macrophages, one of the cell types that ferry virus from the lumen of the airways to the lymphatic organs. To generate macrophages, we isolated monocytes from donated human blood and treated them with the appropriate cytokines to stimulate their differentiation into M2 macrophages (**Fig 4A**). We then co-cultured these M2 macrophages with extruded infectious centers collected from an apical wash of MeV-infected HAE 14 days post-inoculation. As a comparison, we used cell-free virus collected in parallel. Two days later, macrophages were examined for signs of infection using microscopy (**Fig 4B**). Cell-associated virus (green arrow) spread MeV to nearby macrophages (red arrows); cell-free MeV also infected macrophages, but its lower titers limited the effective MOI. When the number of infected macrophages were quantified by visual counting and normalized to the input titer determined post-hoc, we observed similar levels of infectivity between cell-associated and cell-free MeV (**Fig 4C**). These experiments suggest that when normalized to input PFU, infectious centers are as effective as cell-free virus in delivering MeV to macrophages. However, since most virus remains cell-associated, sloughed infectious centers may be the primary infection spreader. We next sought to better understand the mechanism of infectious center release from the epithelial sheet.

**Fig 4.**
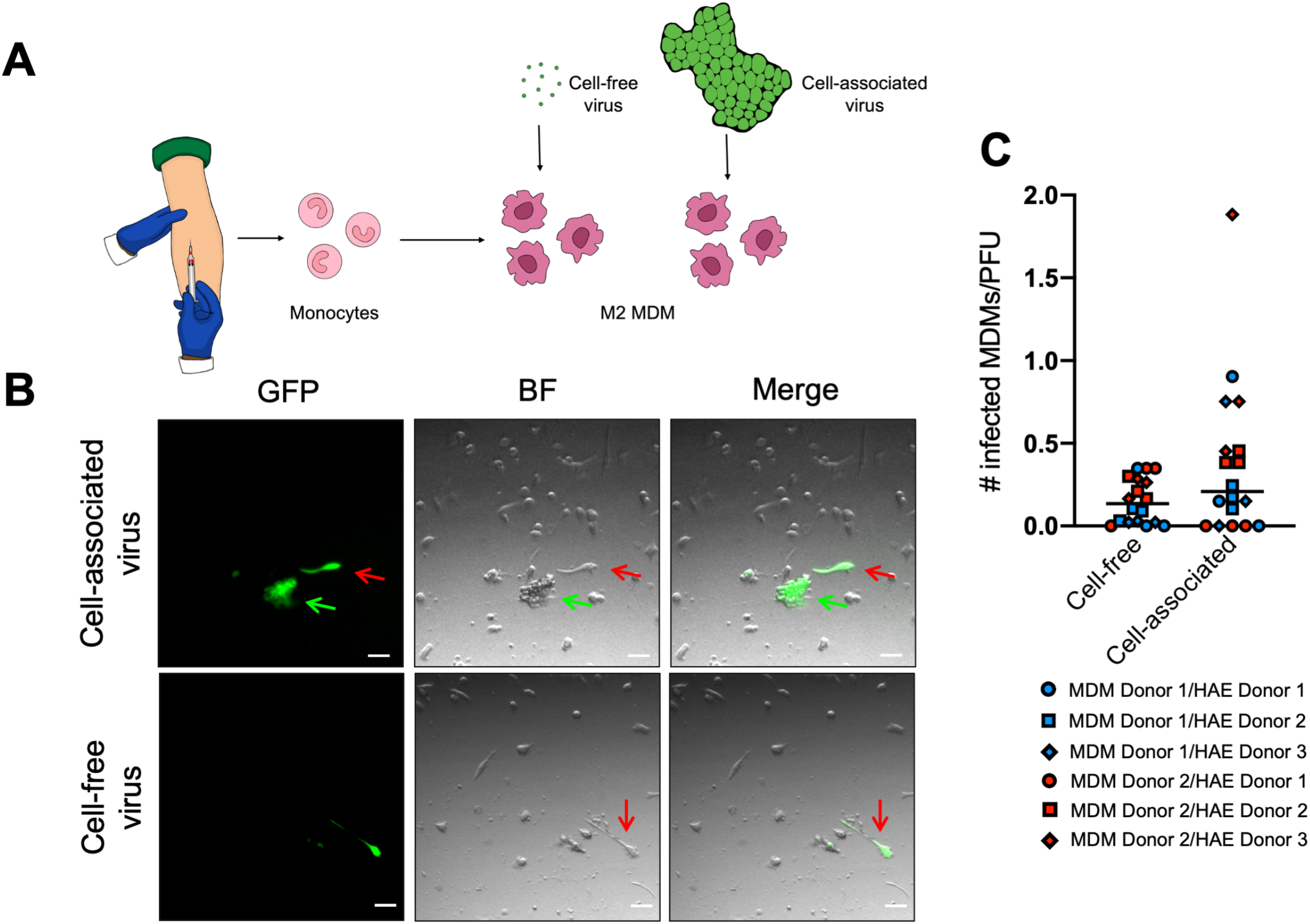
Sloughed infectious centers spread MeV infection. (A) The experimental design is shown schematically. Monocytes were isolated from human donor blood (n = 2 donors) and treated with selected cytokines to induce differentiation into M2 MDMs. Cell-free and cell-associated virus from MeV-infected HAE (14 days post-infection; n = 3) were applied to the macrophages. (B) Spread was evaluated via inverted fluorescent microscopy two days after transfer to macrophages (scale bars = 50 μm; green arrow, cell-associated virus; red arrow, infected macrophages). Images are representative of 3 independent experiments. (C) The cell-associated and cell-free virus was titered concurrently via TCID_50_. Counts of infected macrophages were adjusted for titer differences. A Student’s t-test indicated no statistical significance. MDM, monocyte-derived macrophages; BF, brightfield.

### Defining the transcriptome of MeV infected HAE

To better understand the cellular response to MeV infection, we performed single-cell RNA-seq (scRNA-seq) on infected HAE cultures at 3, 7, and 14 days post-inoculation, and as control, mock-infected HAE at days 3 and 14 (**Fig 5A**). Each condition included cultures from 10 pooled matched human donors; similar numbers of cells were sequenced and subjected to equally powered bioinformatic analyses. In total, RNAs from 30,743 cells were sequenced via 10x Genomics scRNA-seq.

**Fig 5.**
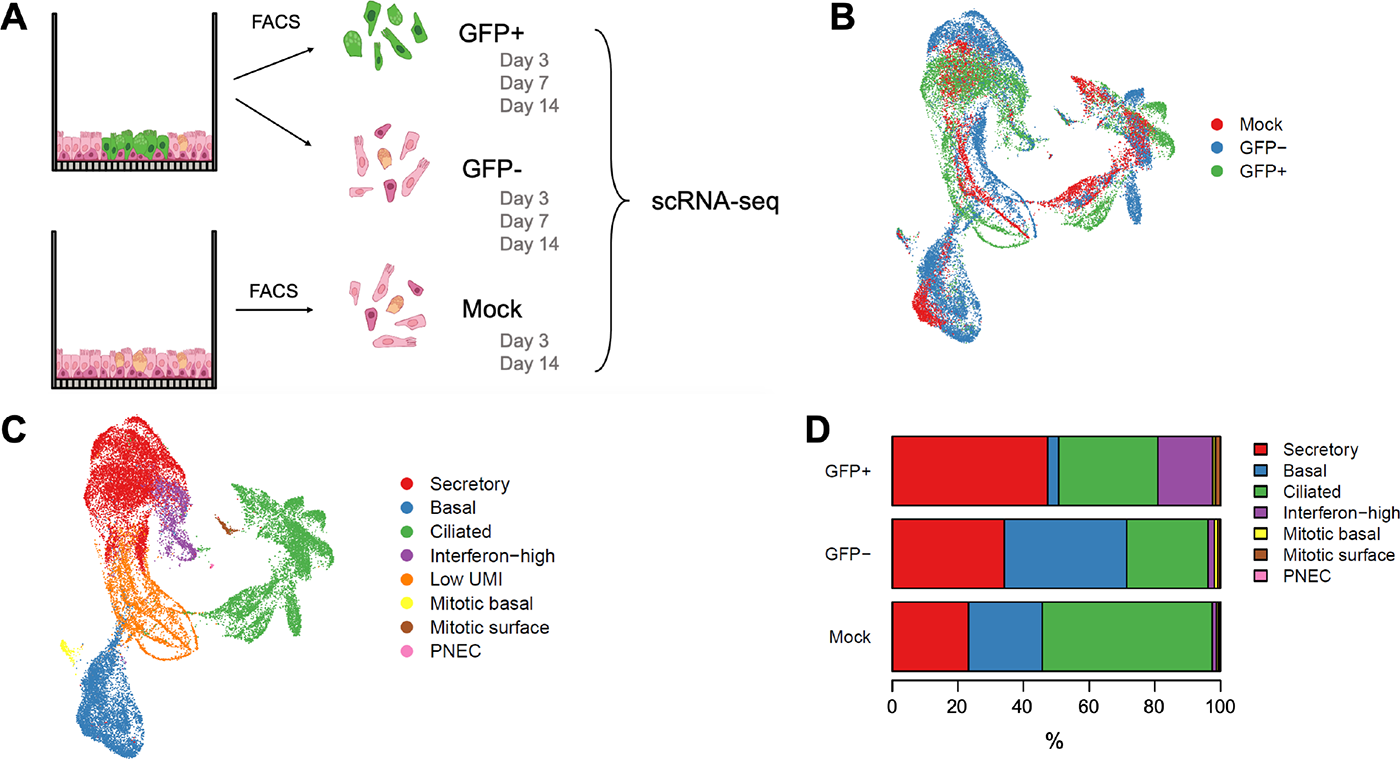
Defining the transcriptome of MeV infected HAE with scRNA-seq. (A) The experimental design is shown schematically. MeV or mock-infected HAE (n = 10 human donors; MOI = 5) were sorted via FACS at day 3, 7, or 14 post-infection and gated for GFP expression. GFP+ and GFP-cells were collected from MeV-infected HAE. Control cells were sorted via FACS from mock infected cultures (referred to as Mock). Cells from all 10 donors were pooled within their treatment type and prepared for scRNA-seq (10x Genomics). In total, 30,743 cells were sequenced. We projected these cells in a Uniform Manifold Approximation and Projection for Dimension Reduction (UMAP) and color-coded them by their treatment group (B) and cell type (C). (D) The percentage of each cell type within each treatment group is shown.

Results were visualized in a uniform manifold approximation and projection (UMAP), where cells with similar gene expression profiles cluster (**Figs 5B, 5C, and S2A**). Similar profile distributions were observed for GFP+, GFP-, and mock-infected cells (**Fig 5B**). Using expression profiles of marker genes (**Fig S2B**), we defined 8 individual clusters (**Fig 5C**), four of them representing the main HAE cell types: secretory, basal, ciliated, and the rare (<1%) pulmonary neuroendocrine cells (PNECs). The four additional clusters were defined by a combination of cell type and phenotypic markers: interferon-high, low unique molecular identifier (UMI), mitotic basal, and mitotic surface.

HAE are typically mitotically quiescent. However, we identified two small, but distinct clusters of dividing cells, mitotic basal and mitotic surface. These clusters are primarily composed of both GFP+ and GFP-cells from the day 14 timepoint in infected cultures and are almost absent in mock-infected cells (**Figs 5C and S2A**). Consistent with microscopic evidence showing that basal cells are non-permissive to MeV infection, GFP+ basal cells were uncommon (**Fig 5D**). Of note, the cell type specificity of interferon-high cells could not be determined, but these cells were predominately GFP+ (**Fig 5C, D**).

We also compared the levels of viral RNAs (vRNAs) for each cell type in infected (GFP+ and GFP-combined) and mock-infected cultures over time (**Fig S2C**). Consistent with earlier observations, vRNA was consistently low in non-dividing basal cells. New observations included the existence of increasing vRNA levels in mitotic basal cells, and high levels of vRNA expression in the newly defined interferon-high cluster at 14 days post-infection.

### Candidate gene expression pathways involved in infectious center sloughing

To identify enriched or reduced biological processes resulting from MeV infection, we performed unbiased signal pathway analysis. As a comparison between the GFP+, GFP-, and Mock groups, lists of differentially expressed genes were generated with a threshold adjusted p-value of 0.05. For GFP+ cells, we identified 91 upregulated genes and 83 downregulated genes; for GFP-cells, 66 upregulated genes and 34 downregulated genes were identified (**Supplemental Table 1**).

A gene ontology analysis tool, GenCLiP 2.0, was then used to identify gene expression pathways activated or repressed during infectious center sloughing [29, 30]. Interferon and inflammation related pathways were more upregulated in GFP+ cells as compared to GFP-cells **(Fig 6A, B)**. Of note, apoptosis pathways were downregulated in GFP+ cells (**Fig 6 A, C**), consistent with our earlier observations (**Fig 3**). In addition, pathways associated with wound healing, cell growth, and cell differentiation were upregulated in GFP-cells as compared to GFP+ cells **(Fig 6 A, B)**. Cell proliferation genes were upregulated in GFP-cells as compared to GFP+ cells throughout the course of infection (**Fig 6D**). Altogether, these results indicate that GFP+ cells inhibit apoptotic pathways as innate immune responses develop. In contrast, GFP-cells begin to differentiate. This suggested to us that basal cells situated underneath infectious centers may divide, possibly in preparation for taking over the epithelium-sealing function of the sloughing infected patch.

**Fig 6.**
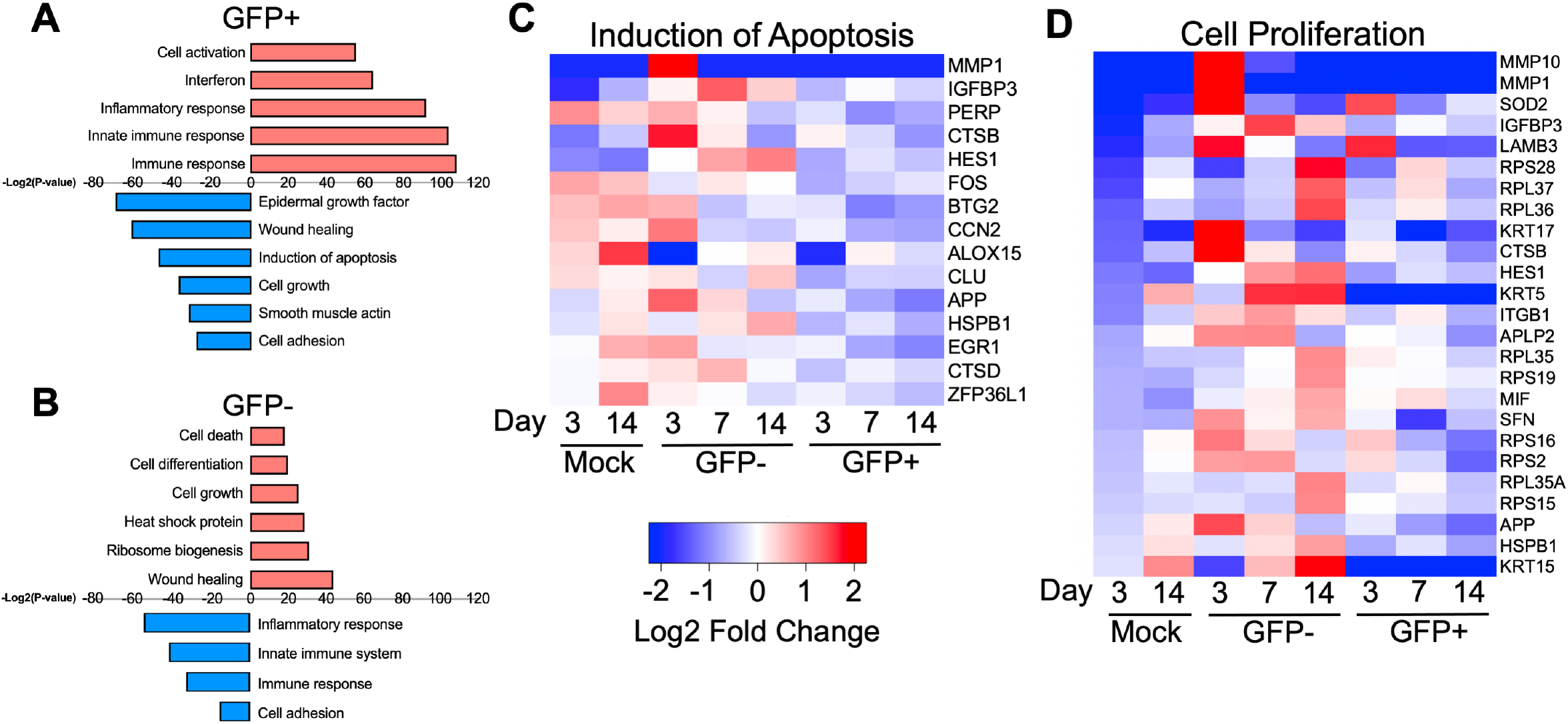
Candidate gene expression pathways involved in infectious center sloughing. Pathway analysis of differentially expressed genes for (A) GFP+ and (B) GFP-cells is shown. Red bars indicate pathways associated with upregulated genes and blue bars indicate pathways associated with downregulated genes. Gene expression heatmaps for genes associated with (C) apoptosis or (D) cell proliferation is shown.

### Basal cells underneath infectious centers proliferate

After confirming that baseline transepithelial electrical resistance remained constant following MeV-GFP infection of HAE (**Fig 7A**), we asked whether cells situated underneath infectious centers proliferate. To identify dividing cells, we used the DNA synthesis marker 5-ethynyl-2’-deoxyuridine (EdU). Indeed, EdU+ cells were localized with infectious centers (**Fig 7B, C**). We then quantified the kinetics of cell division induction below infectious centers. At day 3 post-inoculation, few EdU+ cells were detected in association with infectious centers, but the number of EdU+ cells continuously increased with time (**Fig 7D**). Consistent with this observation, the scRNA-seq dataset indicated an increase of mitotic basal cells over time (**Fig 7E**). A control EdU+ cell count that excluded infectious centers confirmed the quiescent state of cells not located below infectious centers (**Fig 7F, G**). These data show that basal cell proliferation is associated with infectious center formation in HAE. Such proliferation may protect the integrity of the epithelium as infectious centers passively slough or actively dislodge the infectious center from the epithelial sheet.

**Fig 7.**
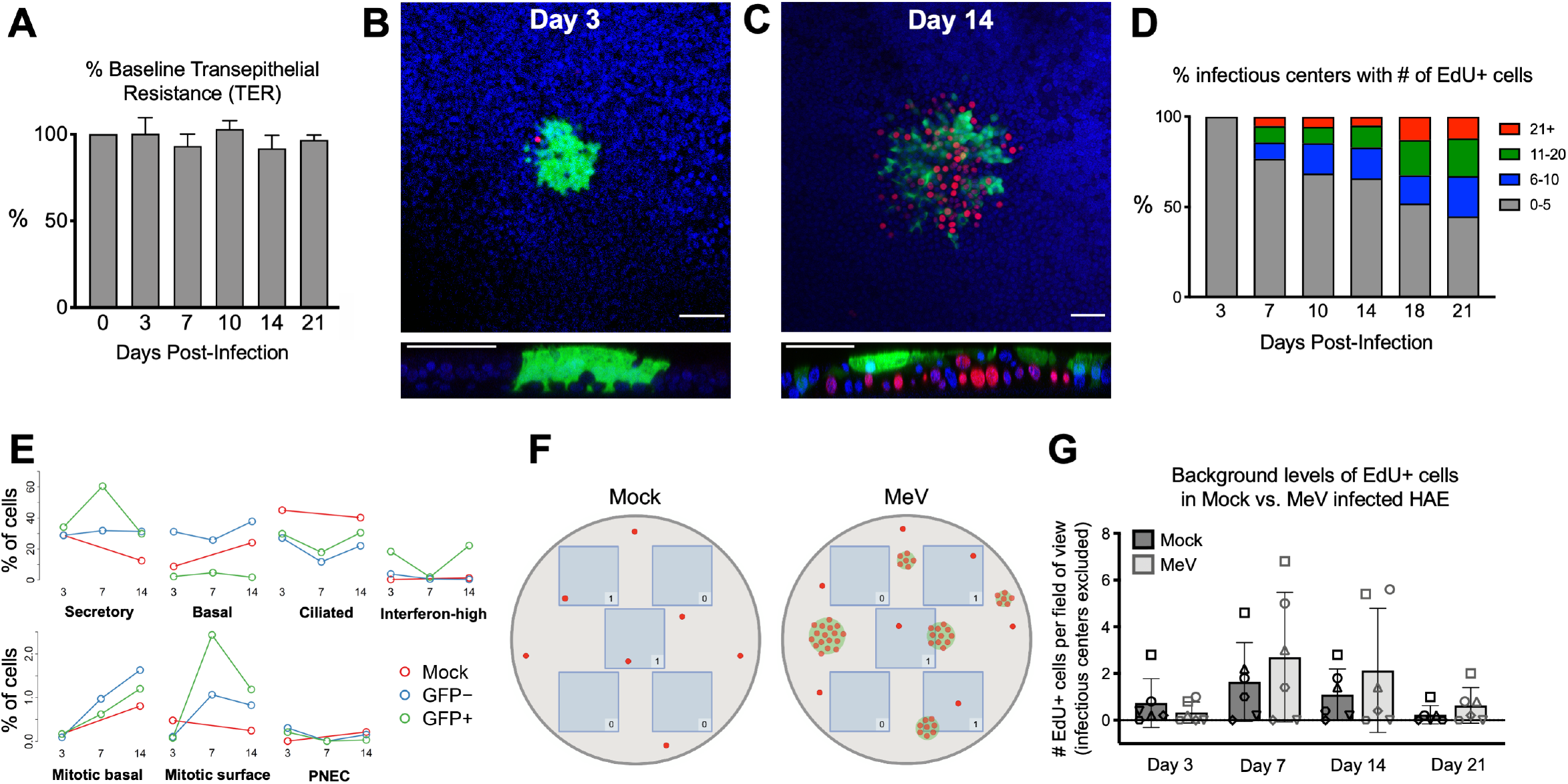
Basal cell proliferation is stimulated underneath infectious centers. (A) Transepithelial electrical resistance (TER) of MeV-infected HAE (MOI = 1; n = 3) over 21 days of infection. Measured by an epithelial ohm meter with a chopstick electrode (EVOM^2^; World Precision Instruments) and shown as a percentage of baseline. EdU immunostaining of MeV-infected HAE at (B) 3 days post-infection and (C) 14 days post-infection. EdU was applied for 16 hours at 10 μM before fixation and staining. Images are representative of 4 independent experiments and 9 human donors. Red, EdU; green, MeV-GFP; blue, DAPI; scale bars = 50 μm. (D) The number of infectious centers and EdU+ cells associated with each infectious center were counted. The key shows the number of EdU+ cells associated with an infectious center. The Y-axis signifies the percentage of infectious centers with that number of EdU+ cells for each timepoint. (E) The percentage of cells identified by scRNA-seq over the time course. (F) A schematic describing the quantification of background proliferation is shown. For each HAE culture (mock or MeV-infected), 5 fields of view were imaged at 20x, identified as blue boxes. All EdU+ cells that fell within the field of view were counted unless they were associated with an infectious center, as represented by the counts in the corner of each box. (G) Quantification was performed on HAE infected for 3, 7, 14, or 21 days (MOI = 1; n = 6 human donors as indicated by a unique shape).

## DISCUSSION

We demonstrate that MeV exits the epithelial sheet via dislodging of infectious centers. We also show that sloughed infectious centers can transmit infection to human macrophages, one of the cell types that carries infectivity from the lumen of a new host’s airways to its lymphatic organs. Since extruded infectious centers contain the most released virus, they may have a central role in host-to-host transmission. Our results also indicate that little cell-free virus is released from HAE, which challenges the idea that apical budding is the major pathway by which MeV exits the airways [31-34].

Infectious center sloughing is consistent with published *in vivo* observations. The presence of exfoliated giant epithelial cells in swab samples from patients is a diagnostic feature of measles [35, 36]. Giant cells can be detected in nasopharyngeal mucus from the start of the measles rash, and the duration of their excretion correlates with severity of acute disease [36]. In bronchial alveolar lavages from experimentally infected macaques, Ludlow et al. documented high numbers of MeV-infected cells or cell debris “spilling” from epithelia into the respiratory tract [25]. These authors also measured equivalent titers of expelled cell-free and cell-associated virus released into the airways and attributed the presence of cells in the airways to stimulation of the cough response. Our data show that coughing is not required to release infected cells. Consistent with *in vivo* observations [25], infectious center sloughing may promote coughing and sneezing that contributes to the infectious nature of MeV. Although the primary spread of MeV appears to be through aerosols and respiratory droplets, fomites coated with sloughed infectious centers could also be a significant contributor in viral transmission [18, 26, 37, 38].

MeV is the most contagious human virus [2-4]. However, the limited currently available transmission studies do not explain why MeV is so much more transmissible than other respiratory viruses. In fact, experimentally infected non-human primates exhibit low MeV titers in bronchial lavage fluid as compared to other respiratory viruses [11, 24, 25]. We think that metabolically active MeV infectious centers could survive in the environment longer than viral particles, and that sloughing of infectious centers accounts in part for the MeV’s high basic reproduction number. Examples of increased viral stability achieved through packaging include the encapsulation of enteroviruses in vesicles, and baculovirus ocular bodies that are more resistant to heat, desiccation, radiation, and chlorine treatment when compared to free virus [39-41]. Vesicle-cloaked rotaviruses are more infectious than free virions, and it is postulated that virions enclosed in vesicles are protected from degradation by intestinal proteases and/or bile acids [41]. Enteric hepatitis A virus (HAV) membrane-encapsulated virions provide protection against neutralizing antibodies that result in enhanced spread within a host [42]. Another advantage of virus delivery through infectious centers is high titer “*en bloc*” transmission of multiple genomes, which may be required for rapid MeV dissemination in its two ecological niches [43, 44]. Further experiments are required to confirm the survival benefits of MeV remaining cell-associated in the environment.

While apoptosis induces cell sloughing for other respiratory viruses [28, 45, 46], our scRNA-seq data, lack of detection of activated caspases, and the documentation of ciliary beating in sloughed infectious centers indicate that MeV can effectively control apoptosis of well-differentiated HAE. Based on insights from gene ontology analyses, future studies will focus on genes controlling wound healing pathways and cell adhesion processes as potential regulators of the sloughing mechanisms. MeV may take advantage of the host response to actively extrude cells that pose a risk to the integrity to the epithelial sheet. Indeed, live cell extrusion from epithelial sheets can result from multiple stimuli, such as overcrowding, tumor suppression, or invasion by pathogens [47-50]. Our results show that basal cell proliferation occurs directly underneath infectious centers. Cell proliferation may reflect the host’s response to replace sloughing or damaged cells, promoting extrusion by “pushing” infectious centers off the epithelial layer. We observed by microscopy, and confirmed though scRNA-seq, that basal cells are rarely infected by MeV. Since basal cells are the primary proliferative cell type in differentiated epithelia, this could explain how the epithelia can maintain integrity for at least 21 days. The relative expression of nectin-4, the epithelial cellular receptor for MeV [12, 13], in basal cells could account, in part, for their nonpermissivity to MeV infection; however, additional studies are required to determine how basal cells are resistant to MeV.

We acknowledge that this study has limitations. First, all experiments were performed *in vitro*. Unpassaged primary HAE cultures recapitulate the *in vivo* airway surface epithelium in cell type distribution and morphology. However, they do not contain immune cells which contribute to clearing infections from the airways and may impact infectious center growth and/or sloughing. Second, the single cell sequencing experiments necessitated sorting to enrich for MeV infected (ie, GFP+) cells and ensure adequate sampling. As a result, we are aware that cell sorting may have skewed our samples toward cell types that are more easily disassociated into single cell populations. Finally, experiments with HAE do not allow us to test the efficacy of host-to-host spread of cell-associated MeV. To address these limitations, future research should include *in vivo* non-human primate studies.

In summary, our results document that MeV uses a novel mechanism of infectious center dislodging to exit airway epithelia. Cell-associated MeV in sloughed infectious centers may be protected from environmental stressors that promote virion degradation during inter-host transmission. Active expelling of infectious centers into the environment may contribute to the exceptionally high transmission efficiency of MeV.

## MATERIALS AND METHODS

### Ethical statement

The well-differentiated primary cultures of human airway epithelia (HAE) in this study were provided by the University of Iowa *In Vitro* Models and Cell Culture Core using discarded tissue, autopsy, or surgical specimens. No identifiable information was provided and all human subject studies were conducted with approval from the University of Iowa Institutional Review Board.

### Human airway epithelial cells

The University of Iowa *In Vitro* Models and Cell Culture Core cultured and maintained HAE as previously described [51]. Briefly, following enzymatic disassociation of trachea and bronchus epithelia, the cells were seeded onto collagen-coated, polycarbonate transwell inserts (0.4 μm pore size; surface area = 0.33 cm^2^; Corning Costar, Cambridge, MA). HAE were submerged in Ultraser G (USG) medium for 24 hours (37°C and 5% CO_2_) at which point the apical media is removed to encourage polarization and differentiation at an air-liquid interface. The HAE used in these experiments were at least 3 weeks old with a transepithelial electrical resistance > 500 Ω.μm^2^.

### Measles virus production

The MeV-GFP virus used in these experiments is a recombinant MeV derived from the wild-type IC-323 strain. The generation and use of this virus have been previously published [7]. Briefly, Vero-hSLAMF1 cells [52] stably express the human measles receptor SLAMF1 and were cultured in Dulbecco modified Eagle medium (DMEM; Thermo Fisher Scientific) containing 5% newborn calf serum (NCS; Thermo Fisher Scientific) and penicillin-streptomycin (100 mg/mL; Thermo Fisher Scientific). After infection with MeV-GFP, the virus is allowed to propagate for 2-3 days at which point the cells are lysed via three freeze/thaw cycles to release the virus. TCID_50_ titers (with Vero-hSLAMF1 cells) are used to determine the titer of MeV-GFP. The titer of MeV-GFP used in these experiments was ∼10^7^ TCID_50_/mL.

### Infection of HAE

Infection of HAE in these experiments was performed as previously described [21, 22]. Briefly, because MeV enters HAE basolaterally, HAE cultures are inverted and covered with a 50 μL mixture of serum-free medium and MeV-GFP. HAE are incubated for 2-4 hours at 37°C and 5% CO_2_ before the inoculum is removed and the cultures are returned upright. RSV infections were accomplished by delivering a 100 μL mixture of serum-free medium and RSV-GFP to the apical side of HAE. After 2 hours of incubation at 37°C and 5% CO_2_, the inoculum was removed and the HAE were washed with serum-free medium three times.

### Separation of cell-free and cell-associated virus

100 μl of USG medium was applied apically to each transwell of MeV-infected HAE. After 5 minutes of incubation (37°C and 5% CO_2_), the medium was gently pipetted up and down two times before collection. Washes were then centrifuged for 3 minutes at 200 x g. The supernatant, containing cell-free virus, was then transferred to a new tube. The pellet, containing cell-associated virus, was resuspended in 100 μl of USG medium.

### Caspase-3 activity assay

MeV-infected, RSV-infected, staurosporine-treated, or mock-infected HAE were assayed for caspase-3 activity using the EnzChek Caspase-3 Assay Kit #1, Z-DEVD-AMC substrate (catalog no. E13183, Invitrogen) in black, clear bottom 96-well assay plates (catalog no. 3603, Corning Costar). Cells were treated apically with 100 μM staurosporine (catalog no. ab120056; Abcam, Cambridge MA) in PBS for 5 hours. Treatment was removed, cells were washed with PBS, and were immediately fixed or assayed. Fluorescence was measured via a SpectraMax i3x Multi-Mode Microplate Reader (Molecular Devices; San Jose, CA).

### Immunostaining and microscopy

Cells were prepared for immunostaining and confocal microscopy by fixation in 2% paraformaldehyde for 15 minutes, permeabilization in 0.2% Triton X-100 for 1 hour, and blocking in SuperBlock Blocking Buffer (Thermo Fisher Scientific, Waltham, MA). Cleaved caspase-3 was immunostained by incubating HAE with a primary human cleaved caspase-3 (Asp175) antibody (catalog no. MAB835; R&D Systems, Minneapolis, MN, 1:100 in SuperBlock Blocking Buffer) for 1 hour. This was followed up with a 1-hour incubation of an Alexa 568 labeled anti-rabbit secondary antibody (catalog no. A-11036; Invitrogen, Waltham, MA, 1:1000 in SuperBlock Blocking Buffer). To stain for F-actin, HAE were incubated with Phalloidin-Alexa 647 (1:50 in PBS, catalog no. A22287; Thermo Fisher Scientific) for 30 minutes. The filters with the HAE were then cut from the rest of the transwell insert and mounted on glass microscope slides using VECTASHIELD Mounting Medium with DAPI (catalog no. H-1200-10; Vector Laboratories, Inc., Burlingame, CA). Confocal images were acquired using a Leica TCS SP3 confocal microscope (Leica Microsystems, Inc.) with 20x, 40x, and 63x objectives. Images were processed and z-stacks were compiled using ImageJ version 2.1.0. Live-image microscopy was performed using a Leica DMI6000-B inverted microscope (Leica Microsystems, Inc., Buffalo Grove, IL) using a 10x objective.

### EdU staining

10 μM 5-Ethynyl-2’-deoxyuridine (EdU) was added to the basolateral media of HAE for 16 hours. HAE were fixed with 2% paraformaldehyde for 15 minutes. HAE were blocked and permeabilized with 3% BSA in PBS and 0.2% Triton X-100 in PBS. The Click-iT EdU Cell Proliferation Kit (Alexa Fluor 594, Thermo Fisher Scientific) was used to detect EdU+ cells. The HAE were washed and mounted on glass slides with VECTASHIELD Mounting Medium with DAPI. Images were taken using confocal microscope and a 40x objective.

### Isolation of primary human monocyte-derived macrophages

Peripheral blood mononuclear cells (PBMCs) were isolated from healthy human donors by performing a Ficoll-Paque gradient (Thermo Fisher) on whole blood. The PBMCs were then cultured in RPMI 1640 medium (supplemented with 10% fetal bovine serum, 5% penicillin/streptomycin, and 1x non-essential amino acid) and 50 ng/mL of human macrophage colony-stimulating factor (M-CSF, Millipore, Temecula, CA) for 5-6 days at 37°C and 5% CO_2_. The cells were then stimulated with 20 ng/mL of recombinant human IL-4 (Gibco) and 20 ng/mL of recombinant human IL-13 (Sigma-Aldrich, St. Louis, MO) to promote differentiation into M2 macrophages. The cells are considered fully differentiated upon observation of a change in morphology (∼7 days post-collection). For the infection experiments, M2 monocyte-derived macrophages were plated on 96-well plates (catalog no. 3596, Corning Costar) at a density of 20,000 cells/well. Cell-free and cell-associated virus were collected as described above. A portion of each collection was set aside for TCID_50_ titers. 50 μL of either cell-free or cell-associated virus was applied to each well of macrophages. TCID_50_ titer results were used to back-calculate the amount of infectious material applied per well.

### Fluorescence-activated cell sorting (FACS)

Mock and MeV-infected HAE (MOI = 5) were prepared for FACS at 3, 7, and 14 days post-infection. HAE were dissociated by incubation with TrypLE (Gibco) for 30 minutes at 37°C and 5% CO_2_. Dissociated cells were collected and centrifuged at 200 x g for 5 minutes. The TrypLE was aspirated, the cells were resuspended in DMEM/F-12 media (Gibco) and kept on ice (∼4°C). FACS was performed on a BD FACSAria Fusion (BD Biosciences, San Jose, CA) by the University of Iowa Flow Cytometry Core.

### Single-cell RNA sequencing (scRNA-seq)

We generated single-cell RNA sequencing libraries using the Chromium Single Cell Gene Expression v3 kit (10X Genomics, Pleasanton, CA). Briefly, ∼5,000 cells from each sample were loaded into a Chromium Next GEM Chip with Gel Beads and Master Mix where they were partitioned in oil to form gel beads in emulsion (GEMs). The GEMs were then barcoded with an Illumina TruSeq sequencing primer, barcode, and unique molecular identifier (UMI). The samples then undergo reverse transcription, cDNA amplification, enzymatic fragmentation, End Repair, A-tailing, Adaptor Ligation, and PCR to finalize the library preparation. The samples were then sequenced by the Genomics Division of the Iowa Institute of Human Genetics using the NovaSeq 6000.

### Bioinformatic analyses

Raw sequencing reads were processed using CellRanger version 3.0.2. Reads were aligned to a hybrid genome consisting of human genome reference GRCh38.p13 and MeV-GFP. Loupe Browser v4.1.0 was used to visualize cells and generate lists of differentially expressed genes. GenClip2.0 was used to identify candidate pathways in a gene ontology analysis. For analysis of gene expression at single cell resolution, gene-by-cell count matrices for each sample were merged and analyzed with the R package Seurat version 3.1.1 [53, 54]. Counts for each cell were normalized by total UMIs and log transformed to quantify gene expression. Centered and scaled gene expression for the 2,000 mostly highly variable genes were reduced to the first 12 principal component scores for input to a shared nearest neighbor clustering algorithm. Cell types were identified by testing for highly upregulated genes in each cluster using a Wilcoxon rank sum test and associating upregulated genes with a list of known airway epithelial markers.

### Western blot

Mock or MeV-infected HAE were lysed using RIPA Lysis and Extraction Buffer (Thermo Fisher Scientific) with complete mini EDTA-free protease inhibitors (Roche, Mannheim, Germany). Protein concentration was determined via the Pierce BCA Protein Assay Kit (Thermo Fisher Scientific). Samples were boiled at 95°C for 5 minutes with Laemmli buffer and 20 μg of each was loaded into a 4-20% Mini-PROTEAN TGX Precast Protein Gel (BioRad, Hercules, CA). Gels were run at 100 V for 30-60 minutes and then transferred to PVDF membranes for 2 hours at 250 mV. Blots were blocked with 5% milk in 1x TBS-T buffer for 1 hour. Primary antibodies for cleaved caspase-3 (catalog no. MAB835; R&D Systems) and polyclonal rabbit anti-N_505_ [55] were used at a concentration of 1:800 and 1:1000 respectively. Horseradish peroxidase (HRP)-conjugated goat anti-rabbit IgG(H+L) (catalog no. 111-035-144, Millipore) was used as a secondary at 1:10,000. Blots were developed with SuperSignal West Pico PLUS Chemiluminescent Substrate (Thermo Fisher Scientific).

### Statistics

Unless otherwise indicated, all numerical data presented in bar graphs are shown as the mean ± SE. Statistical analyses were performed using GraphPad Prism software. Two tailed, unpaired Student’s t tests or one-way ANOVA with Tukey’s correction for multiple comparisons assuming equal variance were used to compare experimental groups. p values <0.05 were considered statistically significant (*p < 0.05, ****p < 0.0001).

## ACKNOWLEDGEMENTS

We thank Steven Varga for providing the RSV-GFP. We thank Jennifer Bartlett and Miguel Ortiz Bezara for their critical reading of the manuscript and Ni Li and Guillermo Romano Ibarra for technical help. We acknowledge the support of the University of Iowa Flow Cytometry Core, the Genomics Division of the Iowa Institute of Human Genetics, and the In Vitro Models and Cell Culture Core. This work was supported by the National Institutes of Health R01 AI-132402 (PLS), R01 AI-143791 (RC), and the Cystic Fibrosis Foundation University of Iowa RDP Bioinformatics Core (AAP).

## Author Contributions

Conceptualization, C.E.H., B.K.S., P.L.S.; data curation, C.E.H, B.K.S., A.L.C, A.L.T.; formal analysis, C.E.H., B.K.S., A.L.T.; funding acquisition, A.A.P., R.C., P.L.S.; investigation, C.E.H., B.K.S., A.L.T., A.L.C.; visualization, C.E.H., B.K.S., A.L.T., P.L.S.; writing – original draft preparation, C.E.H., B.K.S.; writing – review & editing, C.E.H., B.K.S., A.L.T., A.L.C., A.A.P., R.C., P.L.S.

